# BIRDMAn: A Bayesian differential abundance framework that enables robust inference of host-microbe associations

**DOI:** 10.1101/2023.01.30.526328

**Authors:** Gibraan Rahman, James T. Morton, Cameron Martino, Gregory D. Sepich-Poore, Celeste Allaband, Caitlin Guccione, Yang Chen, Daniel Hakim, Mehrbod Estaki, Rob Knight

## Abstract

Quantifying the differential abundance (DA) of specific taxa among experimental groups in microbiome studies is challenging due to data characteristics (e.g., compositionality, sparsity) and specific study designs (e.g., repeated measures, meta-analysis, cross-over). Here we present BIRDMAn (**B**ayesian **I**nferential **R**egression for **D**ifferential **M**icrobiome **An**alysis), a flexible DA method that can account for microbiome data characteristics and diverse experimental designs. Simulations show that BIRDMAn models are robust to uneven sequencing depth and provide a >20-fold improvement in statistical power over existing methods. We then use BIRDMAn to identify antibiotic-mediated perturbations undetected by other DA methods due to subject-level heterogeneity. Finally, we demonstrate how BIRDMAn can construct state-of-the-art cancer-type classifiers using The Cancer Genome Atlas (TCGA) dataset, with substantial accuracy improvements over random forests and existing DA tools across multiple sequencing centers. Collectively, BIRDMAn extracts more informative biological signals while accounting for study-specific experimental conditions than existing approaches.

## Main

Advances in sequencing technology and computational methods have enabled researchers to experimentally characterize microbiomes across wide ranges of biological conditions, including psychiatric diseases^1,2^, cancer^3,4^, and COVID-19^5,6^. However, as the understanding of microbial effects on human health and disease has increased, the experimental questions, hypotheses, and concomitant statistics have grown in complexity, with study designs now commonly involving longitudinal analyses^7–9^, experimental interventions^10–12^, and meta-analyses^7^. Although such approaches can provide mechanistic insights into the microbiome’s effect(s) on the host, their conclusions are often limited by the ability to perform valid statistical analyses that are sufficiently flexible to account for the added experimental complexity.

One common but critical challenge in these contexts is when population-level heterogeneity (such as subject-to-subject variation) is confounded by technical variability. For example, samples originating from the same sequencing center will tend to be more similar to each other than those sequenced from different centers^13^. The confounding factors that may explain these differences make it difficult to determine consistent microbial biomarkers associated with biological variables or conditions of interest^8^—an effect compounded by other microbiome data difficulties, such as high sparsity, high-dimensionality, and compositionality. Moreover, statistical tools that can properly assess and account for strong structural effects while still indicating which microbes truly vary between biological conditions are limited to date^15^.

Making matters more difficult, disagreement exists about how to benchmark differential abundance (DA) tools and methods. Previous efforts have commonly focused on comparing the results of hypothesis testing while accounting for the multiplicity of features through false-discovery-rate (FDR) correction^15–17^. Studies have demonstrated that tools designed for differential abundance often report contradictory results with different microbial abundances among biologically distinct sampling groups^19^.

Addressing these challenges requires a more robust statistical framework for benchmarking differential abundance methods and would benefit from flexible DA modeling approaches. Thus, we developed BIRDMAn (**B**ayesian **I**nferential **R**egression for **D**ifferential **M**icrobiome **An**alysis), a flexible computational framework for hierarchical Bayesian modeling of microbiome data that simultaneously accounts for its high sparsity, high-dimensionality, and compositionality.

The Bayesian approach to statistical modeling provides unique advantages compared to frequentist solutions, such as the inclusion of prior information, uncertainty estimation of parameters, native hierarchical modeling, and edge case smoothing (e.g., estimating log fold changes when a feature is only present in one group). Implemented within the Stan programming language (commonly used for designing probabilistic models), BIRDMAn flexibly enables parameter estimation of all biological variables and non-biological covariates. These advantages allow us to demonstrate how explicitly modeling population-level effects in probabilistic BIRDMAn models increases the amount of true biological signal recovered compared to existing tools on both simulated and real-world datasets. Moreover, the BIRDMAn workflow significantly lowers the barrier of entry for differential abundance methods development and implementation. Additionally, to address reproducibility issues of prior DA tool benchmarking, we present a novel approach that employs techniques from compositional data analysis, making the comparison of tools more interpretable and statistically valid.

## Results

BIRDMAn is implemented as a Python interface to the Stan probabilistic programming language, which utilizes Hamiltonian Monte Carlo sampling, one of the state-of-the-art approaches for Bayesian uncertainty estimation^20^. Users can employ pre-configured model designs or flexibly customize inputs to account for their specific experimental design and biological questions; BIRDMAn then fits and processes these models (Fig 1). The results of these analyses are the posterior distributions of the defined parameters of interest, such as log-fold changes and their uncertainty given the data (see Methods).

**Fig 1:**
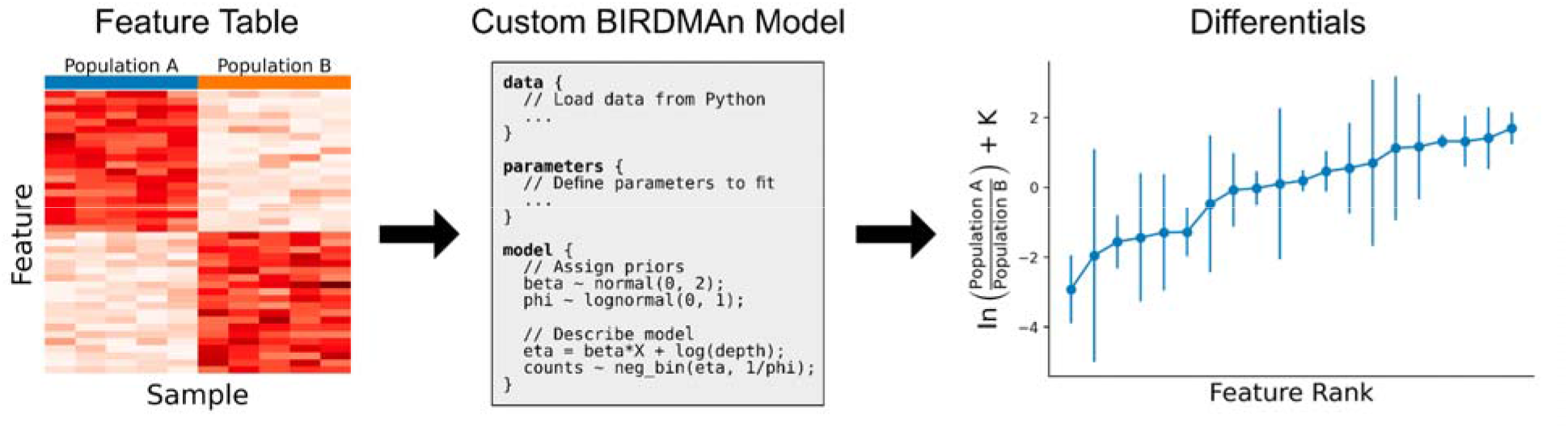
Overview of BIRDMAn workflow for customizable differential abundance analysis. A table of counts by features is modeled using Bayesian probabilistic programming, resulting in credible intervals of the estimated parameter posterior distributions. The statistical model can be customized using the Stan probabilistic programming language and fit using the BIRDMAn Python interface.

To showcase the statistical properties of BIRDMAn models, we first leverage simulations to evaluate the accuracy of estimating differential uncertainty in the context of realistic biological scenarios. Then, we apply BIRDMAn models on real-world data, demonstrating superiority for resolving subject-level heterogeneity in an antibiotics experiment, as well as alleviating sequencing center-specific effects in a cancer genomics dataset, each while capturing biologically-informative signals.

### Simulations demonstrate BIRDMAn model accuracy and precision

A common difficulty in benchmarking differential abundance methods is the lack of ground truth. We typically do not know which microbial taxa are truly increasing or decreasing across experimental conditions. To gain insights into the robustness of BIRDMAn models, we performed a data-driven simulation of a case-control microbiome dataset with one binary covariate, large batch effects (10 features, 10 batches, and 300 samples), data overdispersion, and known differentials associated with case status (see Methods) (Fig 2a). We then used BIRDMAn to estimate the model parameters for each feature and compared the Bayesian posterior estimates with the true value, finding that BIRDMAn models recovered the ground truth differentials with high accuracy and precision (Fig 2b) while outperforming other tools in terms of root mean square error (RMSE) (Fig 2c). This highlights how BIRDMAn model customization permits more accurate estimations of differentials.

**Fig 2:**
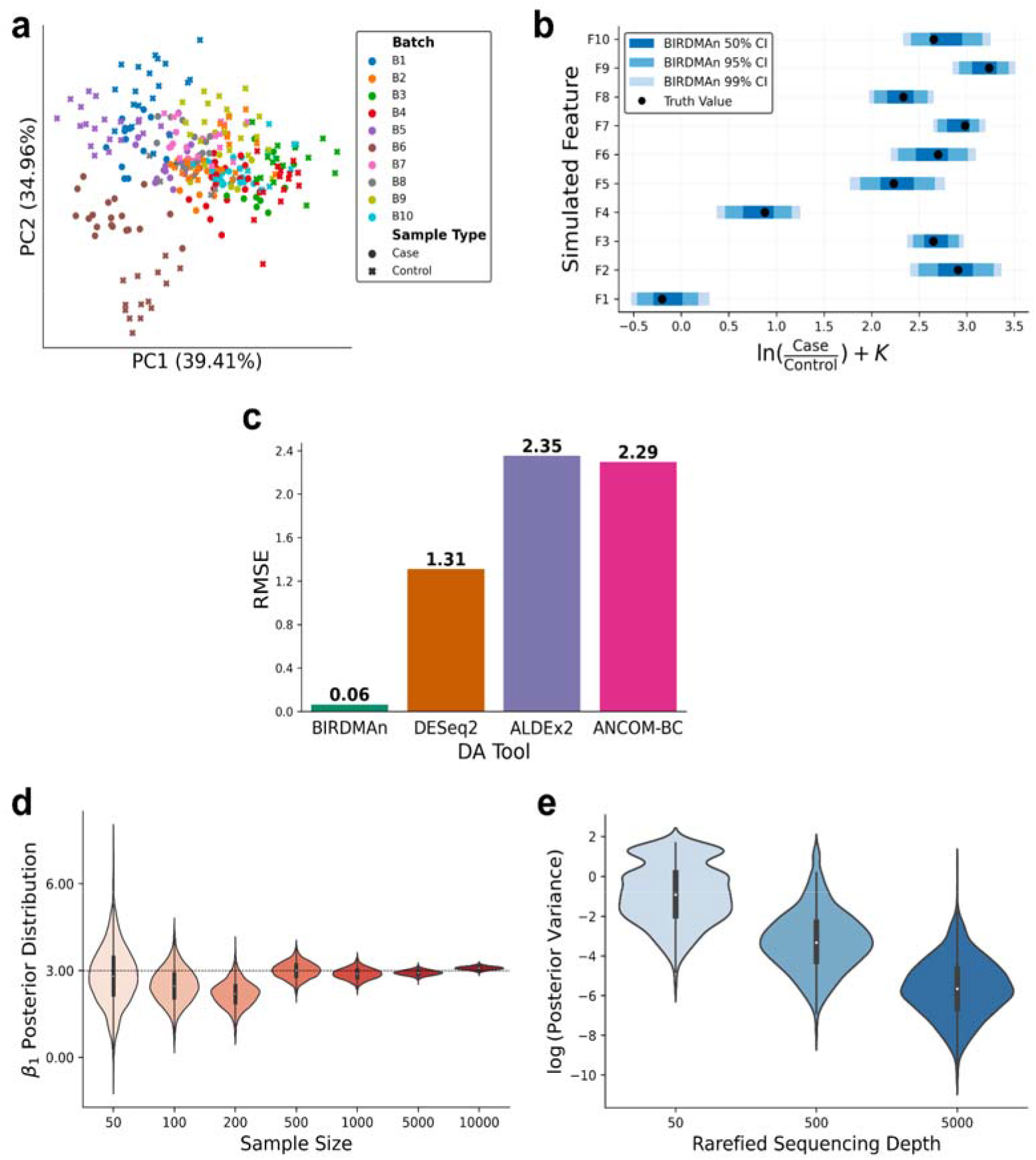
(a) Robust Aitchison principal components plot of the simulated data, showing the large separation by batch effect. Simulations of 10 batches (B1 to B10) of microbiome results, each containing 10 features (F1 to F10), where each feature has a true differential abundance between cases and controls that is the same for each batch, and also a random per batch bias. (b) Recovery of the true simulated log ratio between cases and controls for each feature (black dots), with credible intervals on average centered on the true log ratio (blue bars). (c) Superior performance of BIRDMAn over other differential abundance methods in minimizing the RMSE of the difference between the estimated mean posterior log ratio between cases and controls, revealing a >20-fold improvement in RMSE over the nearest competitor, DESeq2. (d) Estimated distributions of log-fold changes from Bayesian analysis tighten as the number of samples increases. Dashed line represents the true simulated value for each simulation. (e) Rarefaction simulation performed using multinomial count generative models (1000 features) at three different sequencing depths shows that the variance of the posterior distribution decreases as depth increases.

One advantage of Bayesian models is that they can leverage posterior estimates to summarize the uncertainty of these differentials, taking into account the sample size and the sequencing depth. As expected, we show that when BIRDMAn models are fitted on larger sample sizes, the uncertainty decreases, highlighting how incorporating more data, and avoiding rarefaction, enables a more accurate estimation of the differentials (Fig 2d). Furthermore, we show that decreasing the sequencing depth also increases the uncertainty, highlighting how rarefaction could degrade parameter estimates’ precisions in BIRDMAn models (Fig 2e). Since BIRDMAn can handle variable sequencing depths, there is no need to perform rarefaction before model fitting, which is desirable when analyzing microbiome datasets^21^.

### BIRDMAn models capture biological signals missed by other methods during dual-course longitudinal antibiotics

Another challenge for DA methods is to compare multiple samples from the same subject longitudinally (repeated measures) since concomitant host-specific variation can obscure phenotypically-associated microbial changes. Methods designed for longitudinal data^22–26^ cannot easily account for modeling perturbations and struggle with scaling to high dimensions. To demonstrate the use of BIRDMAn on repeated measure study designs, we evaluated a published longitudinal study of two courses of the antibiotic ciprofloxacin (Cp) (3 subjects, 7 timepoints)^27^. Notably, this study originally concluded that inter-subject variability drove the response to antibiotics by examining beta-diversities, which do not account for auto-correlation effects of repeated measures^28^ (Fig 3a). Other studies have also highlighted the importance of properly accounting for the microbial community composition prior to antibiotics when assessing varying responses^29,30^, which requires accurate temporal modeling.

**Fig 3:**
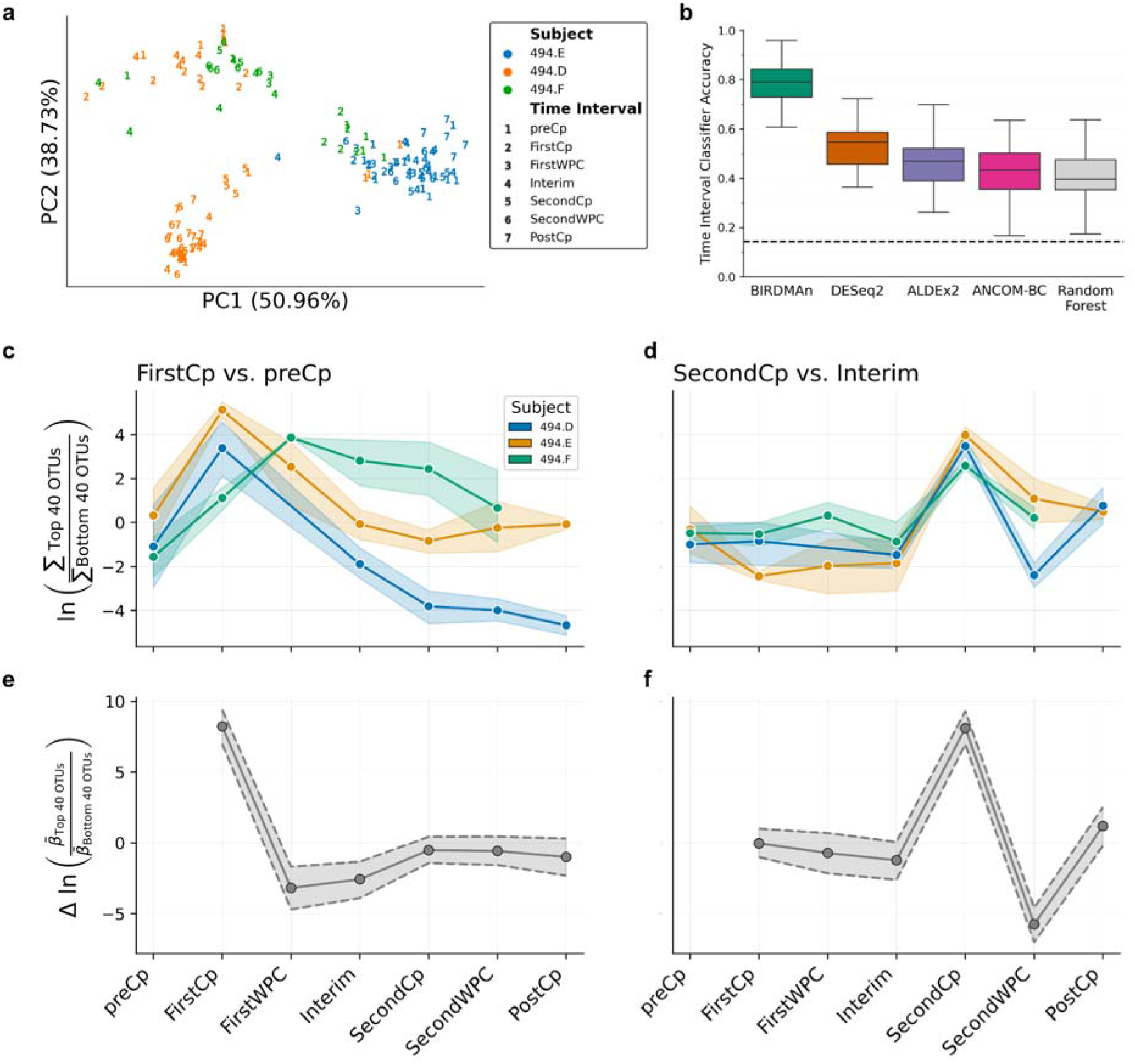
(a) Robust Aitchison principal components plot of full dataset shows samples cluster primarily by host subject. (b) Balanced accuracy of multinomial classification of time point by tool. Differential abundance classifiers were constructed using logistic regression with the log-ratios of the top 40 and bottom 40 OTUs associated with each timepoint as predictors. Repeated k-fold cross-validation was performed with 5 splits and 10 repeats. The mean classifier error is at least twice as great with all other differential abundance tools as with BIRDMAn. Dashed line represents random guessing performance among the seven timepoints. (c, d) Dynamics of sample log-ratios of (c) first Cp course and (d) second Cp course colored by subject. (e, f) Dynamics of BIRDMAn-estimated log-fold changes associated with (e) FirstCp effect with preCp as reference and (f) SecondCp effect with Interim as reference. Shaded intervals represent the 90% credible interval of the estimated posterior distributions.

Given BIRDMAn’s flexibility, we constructed a customized DA model that leverages Linear Mixed Effects models, accounting for repeated measurements from subjects while computing temporal differences (see Methods). This model design then enabled the exploration of common microbial community changes associated with antibiotic perturbation, which the originally published methods could not identify. With the computed log-fold changes over time (Supp Fig 1a), we investigated how consistent antibiotic induced shifts were across subjects. For each temporal difference, we took the top and bottom 40 OTUs to calculate sample log-ratios, which were used to predict antibiotics intake^31^. From these log-ratios, we observed strong, statistically significant temporal shifts associated with each successive time interval (Supp Fig 1b).

To determine if existing tools could have identified these timepoint-specific perturbations, we also developed a multinomial logistic regression classifier based on the BIRDMAn results to predict the corresponding time interval. We then compared our prediction performances against classifiers built using ALDEx2^32^, ANCOM-BC^33^, and DESeq2^34^ results on the same samples, as well as a classifier built on the center log-ratio transformed table (see Methods). Remarkably, BIRDMAn-informed classifiers were able to accurately differentiate between the different treatment groups (accuracy > 0.65) (Supp Fig 1c) and showed substantially better prediction accuracy compared to all other methods (Fig 3b). We also verified that this superior performance held across varying numbers of OTUs used in log-ratio calculation (Supp Fig 1d). Ultimately, these findings show how BIRDMAn can identify clear-cut biological changes that were missed or obscured by other approaches, highlighting its ability to confirm expected biological hypotheses.

We used the sample log-ratios associated with the First and Second Cp applications and plotted the dynamics over time (Fig 3c, d). Accordingly, we plotted the corresponding derivative log-fold changes computed from BIRDMAn (Fig 3e, f) and see that our trajectories match between the sample log-ratios and the estimated log-fold changes, indicating that our model was able to successfully capture the overall signal independent of subject.

The antibiotic used in the original work, Cp, is known to primarily target (though not exclusively) gram negative bacteria^35,36^. We thus hypothesized that the differential abundance results should reflect the longitudinal dynamics of gram negative bacterial abundance. In the top and bottom 40 most changed taxa after FirstCp, 17.5% of the numerator taxa were gram negative, whereas 27.5% of the denominator were gram negative (Supp Fig 2e). Given the Cp antibiotic mechanism, it is likely that gram negative taxa in the denominator decreased which caused the increased log-ratio^37,38^ (Figure 2c). We see that there is a sharp decrease in this log-ratio at FirstWPC, which could be attributed to gut homeostasis^37,38^. However, we see a weaker pattern in the top/bottom 40 microbes after SecondCp, where 2.5% of the numerator taxa were gram negative and 10% of the denominator taxa were gram negative. In contrast to the FirstCp, the microbes most affected by SecondCp quickly returned to their original abundances. Furthermore, we see that the microbes most altered by FirstCp were not affected by SecondCp. Altogether this hints at newly acquired antimicrobial resistant genes after the application of FirstCp.

### BIRDMAn models mitigate batch effects in cancer microbiome data

To investigate how generalizable BIRDMAn models are with respect to population heterogeneity, we conducted a meta-analysis using cancer microbiome data derived from The Cancer Genome Atlas (TCGA). This dataset is known to have large structural batch effects^4^, where the samples were processed at multiple centers across North America, resulting in an artificial separation of cancer microbiomes by sequencing center if not otherwise accounted for (Fig 4a, Supp Fig 2a)^4,39^. These effects can make it difficult to determine microbial biomarkers associated with tumors rather than artifacts of technical variation, but correcting for this could enable downstream host-microbial cancer analyses. We thus tested how well BIRDMAn models could extract biological signals from this dataset while accounting for technical batch effects modeled as random effects. We additionally modeled each microbial feature’s abundance using this approach to determine the specificity of these microbes for each cancer type (see Methods and Code).

**Fig 4:**
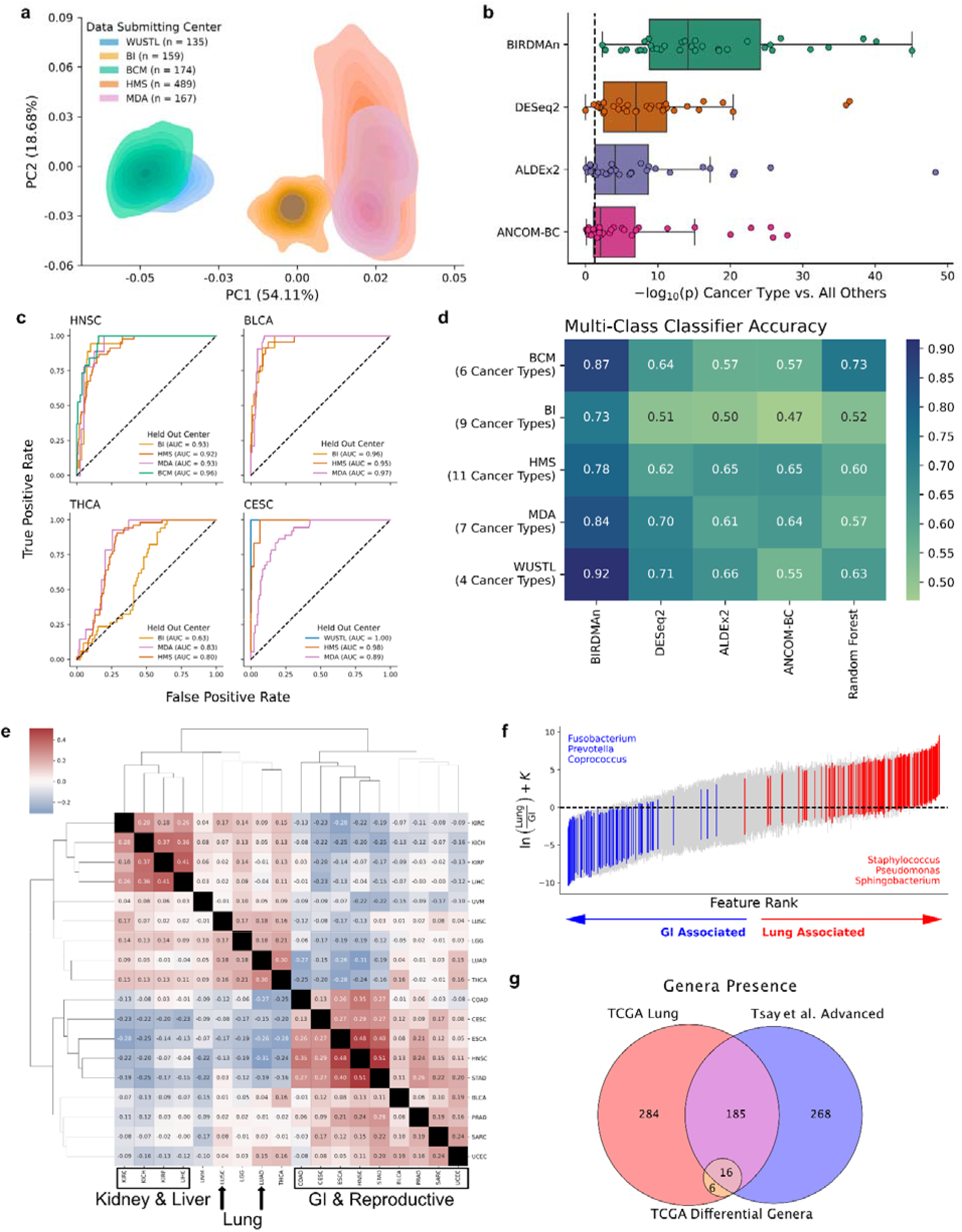
(a) Whole-genome sequenced cancer microbiome data from TCGA shows strong batch effects by sequencing center (colored by center; see Supp Fig 2a for per cancer type plots). Samples are summarized by the 2D kernel density estimate for each center. (b) T-test p-values comparing log-ratios of each cancer type vs. all others within each center. Dashed line represents p=0.05. All differential abundance methods show significant differences with log-ratios to separate the microbes in each individual cancer type from those found in all other cancer types, but BIRDMAn outperforms other methods in highlighting this difference. (c) ROC curves for leave-one-center-out cross-validation for four cancer types where at least 3 centers sequenced that cancer type (BRCA was not included as it was used as reference). Classifiers were built to predict one-vs-rest for that cancer type. BI = Broad Institute of MIT and Harvard; BCM = Baylor College of Medicine; HMS = Harvard Medical School; MDA = MD Anderson Institute for Applied Cancer Science; WUSTL = Washington University School of Medicine. (d) Multinomial (mean) classification accuracy of classifiers to predict cancer type given the log-ratios computed from the top and bottom 200 taxa associated with each cancer type. Random Forests classifier, which is frequently used in this field but is not based on differential abundance, was included as a comparison for this class of methods. Classifications were performed within each center to remove batch effects from predictions. BIRDMAn outperforms all other methods, including Random Forests, for all tumor types. (e) Clustermap of Kendall tau correlation coefficients of pairwise cancer type differentials (breast cancer as reference). (f) Comparison of lung-associated genera with GI-associated genera. Highlighted genera are known to be associated with either lung or GI microbiome and show strong directionality in the BIRDMAn results. (g) Venn diagram of genera present in TCGA lung samples and genera present in advanced stage lung cancer from work published by Tsay et al. Additionally, the 22 genera represented in the top 100 features associated with TCGA lung cancer cancers are included. A majority of these genera (16/22) are present in both datasets.

Since cancer types are known to have distinct microbiomes^4,40^, we first confirmed that BIRDMAn models could extract cancer type-specific differences despite the technical variation observed in this study. From our log-ratio classification benchmarks, we observe that our custom BIRDMAn model can detect a substantially stronger differential signature between the cancer types compared to ALDEx2, ANCOM-BC, DESeq2, and Random Forests (Fig 4b; note the axis log-scaling) after controlling for the batch effects due to the sequencing center (Supp Fig 2c).

To determine the generalizability of our results, we then constructed a leave-one-center-out cross-validation benchmark using logistic regression on the BIRDMAn-computed log-ratios. Four cancer types with at least three represented data submitting centers (head and neck cancer [HNSC], bladder cancer [BLCA], thyroid cancer [THCA], and cervical cancer [CESC]) were included in this benchmark. The receiver operating characteristic (ROC) curves demonstrated strong classification performance (Fig 4c), indicating that BIRDMAn captures generalizable microbial signals across multiple sequencing centers. Generalizability can be a major challenge in microbiome studies^3^, where classifiers become overfitted for individual cohorts. We observe this with other DA tools (ALDEx2, DESeq2, ANCOM-BC) and even Random Forests (Supp Fig 2d), where most tools struggle to achieve an area under the ROC curves (AUROC) of >0.8. BIRDMAn is competitive with these tools, achieving an AUROC >0.9 in HNSC, BLCA, and CESC cancers while achieving the highest predictive accuracy in BLCA and CESC cancers. The high classifier accuracy leaving out each individual center demonstrates that no one center’s data strongly affects the classifier accuracy, with the exception of BI for THCA.

To investigate the heterogeneity across different cancer types, we next computed Kendall correlations of BIRDMAn-estimated microbial log-fold changes across all pairs of cancer types. This analysis revealed similarities among cancer types that we would expect, including strong similarities between kidney cancer subtypes (KIRC, KICH, KIRP), lung cancer subtypes (LUAD, LUSC), and gastrointestinal (GI) cancers (COAD, ESCA, HNSC, STAD), Additionally, the BIRDMAn-informed data suggested some novel associations, such as the similarity between kidney cancers and liver cancer (LIHC). When clustering the individual microbes’ differentials (Supp Fig 2b), we also observed that numerous GI-specific microbes differentiated GI cancers from other cancer types.

When focusing on comparing GI cancers to lung cancers, we found that the resulting BIRDMAn log-fold changes accurately reflected known biology surrounding the niches in which these microbiomes are commonly found. Specifically, *Fusobacterium*^41^, *Prevotella*^42^, and *Coproccus*^43^ are genera commonly found in the GI tract; conversely, *Pseudomonas*^44^, *Staphyloccus*^45^, and *Sphingobacterium*^46^ genera include opportunistic pathogens that are commonly found in lung infections (Fig 4f). We cross-referenced our results against the Tsay *et al*. cohort that utilized 16S rRNA sequencing to investigate lung cancer. Out of the 469 genera in the TCGA lung issues, we observed that 39% of these microbes were also observed in the Tsay *et al*. cohort, despite known previous discordant findings comparing 16S rRNA sequencing and whole genome sequencing^47,48^. Furthermore, when we focus on the top 100 microbes that are detected to be associated with lung cancer, 70% of the represented genera were observed in both the TCGA and Tsay *et al*. datasets. Altogether, this shows how BIRDMAn models can provide biologically-informative results while properly accounting for and mitigating strong structural batch effects that currently confound other DA approaches.

## Discussion

Advances in Bayesian computation have lowered the barriers to developing statistical workflows. To empower microbiome scientists to take advantage of these methods, we developed and implemented a novel approach to differential abundance based on Bayesian hierarchical modeling, with advantages highlighted in simulation benchmarks and real-world datasets. Chiefly, BIRDMAn is designed as a *framework* for researchers to account for the statistical constraints specific to their biological questions. We have demonstrated the benefits of this framework in common biological scenarios involving longitudinal study designs and sequencing center variation — where BIRDMAn can better correct for technical variation than existing methods while identifying biologically-relevant signals. In addition to the ability to construct novel DA models, we presented a robust method for benchmarking and comparing results from different DA tools. In contrast to previous efforts investigating FDR in simulation and reproducibility benchmarks^19,49,50^, we show how to construct sample classifiers from the log-fold change estimates, enabling machine learning techniques such as cross-validation on biological datasets.

Another key challenge of DA benchmarking is the absence of “ground truth,” or the true differentials associated with biological conditions, especially in the presence of strong batch effects. Simulations with known parameters for batch and biological effects can address this limitation, and we showed that BIRDMAn models could recover, with high accuracy and precision, these parameters and their uncertainty. Additional simulations on parameter uncertainty further showed decreases with increased sample size and higher sequencing depth, corroborating previous work and traditional statistical knowledge.

We then investigated two real-world case studies—antibiotics response/recovery and cancer microbiome interactions—demonstrating how BIRDMAn can uncover expected and novel biology. For each dataset, BIRDMAn models were able to account for the inherent effects of center/subject on individual microbial abundances while, when necessary, accounting for complex statistical factors (such as, random intercepts, random slopes, overdispersion). To date, there is no other DA tool that provides a similar and necessary degree of flexible statistical modeling. Our results on the previously published antibiotics dataset revealed the attenuating effect of repeated Cp courses on Gram-negative bacteria, with potential implications for clinical practice using antibiotics. Additionally, BIRDMAn-informed results from the cancer microbiome dataset could be useful in developing novel diagnostic and therapeutic strategies that target or perturb cancer-specific features.

In light of our findings, there are notable assumptions that need to be considered. Specifically, the choice of prior distributions affects the estimated posterior distributions, especially at low sample sizes. Although priors allow researchers to include their expertise in their modeling procedure, it is often the case that an appropriate prior distribution is unknown, requiring uninformed priors with high uncertainty to be used. However, we note that as more analyses are performed, their results can provide a rationale for picking future priors—a strong advantage of the Bayesian approach over non-Bayesian methods. For our purposes, we defined the same prior distribution for each feature within a dataset, but this can easily be adapted to better model features with their expected parameter range. We also note that the (common) lack of absolute abundance data is a limitation in evaluating differential abundance^51^. Strategies to account for this, such as in Williamson *et al*.^52^, could potentially be translated into BIRDMAn models to augment the modeling results. Furthermore, we model the microbial abundances using the negative binomial approach, which is currently contested as an appropriate model for sequencing count data^53^. Still, an advantage of BIRDMAn is that the likelihood function is not restricted to the negative binomial, and one can exchange it for the Poisson-Lognormal, Multinomial, or any other count distribution^54^.

To summarize, we find that careful statistical consideration during DA analysis enables the identification of microbe-phenotype associations that are missed by existing tools. The flexibility of BIRDMAn can thoroughly account for unwanted confounding factors, such as batch and subject, resulting in higher confidence in reported microbial biomarkers. Moreover, the presented log-ratio benchmarking approach opens up numerous possibilities for testing improved machine learning capabilities on microbiome data. Overall, we posit that BIRDMAn’s flexibility and utility will provide impactful statistical results for complex study designs while enabling reproducible science in the microbiome field.

## Methods

### Performing Bayesian inference with Stan

Parameter estimation was performed using Bayesian inference. Our approach utilizes Bayes’ Rule where ***θ*** represents the parameter space and ***D*** represents our collected data:

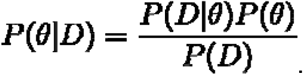

Because the evidence term, ***P(D)*** is simply a normalizing constant, we can rewrite Bayes’ Rule as follows, substituting terms with their common nomenclature:

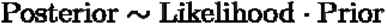

Thus, our objective with Bayesian inference is to obtain the posterior distribution by modeling the likelihood function of our data as well as our prior knowledge of the parameters. Absent a model formulation involving conjugate priors, we cannot compute the posterior distribution analytically. Instead, we use Stan to draw samples from the posterior distribution using the No-U-Turn Hamiltonian Monte Carlo sampler^20^. A series of Markov chains are initialized and allowed to “warm-up” in their exploration of the parameter posterior distributions. Once the defined number of warm-up iterations has concluded, a set number of samples are drawn from each of the chains. Multiple chains are run to ensure that model convergence occurs.

We implement Bayesian inference using the CmdStanPy interface in Python, calling the C++ Stan toolchain for efficient sampling. The warm-up iterations are discarded by default and the sampling iterations are saved for each Markov chain.

### Negative binomial model parameterization

We fit counts of each microbe in a dataset according to a negative binomial distribution as an approximation of multinomial logistic regression^55^. Due to overdispersion, standard count models such as Poisson are inappropriate for sequencing data^21^. We note that the negative binomial model can be considered an extension to the Poisson model with additional variance components^56^.

The negative binomial models used in this work are described by parameters for both mean and overdispersion. This is in contrast to traditional parameters in negative binomial models described by the probability of success and the number of failures before an instance of a success. The former model, often referred to as the “alternative parameterization,” is more amenable to generalized linear modeling through hierarchical models as the mean can be modeled directly.

The basic format of the alternative parameterization negative binomial model is described below where *n* corresponds to the count, *ϕ* the overdispersion, and ***μ*** the mean count.

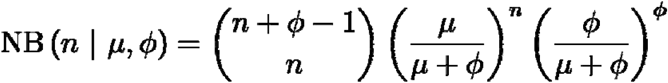

We use a log-link function, ***μ = exp (η)*** to model the mean where the log mean count, ***η***, can be represented by linear terms. To account for variable sequencing depth among samples, we include log sequencing depth as an offset term in our models.

### BIRDMAn framework

We developed BIRDMAn as a framework for highly-customizable Bayesian differential abundance modeling. BIRDMAn abstracts much of the Bayesian workflow away for usage with microbiome data. An object-oriented approach allows users to subclass basic models for their custom implementations. BIRDMAn includes, by default, a Negative Binomial model implementation. This can be used without writing any new Stan code or subclassing any BIRDMAn objects.

BIRDMAn models take BIOM tables^57^ as input containing the sample and observation IDs. Sample metadata can be provided as Pandas DataFrames. We provide a method, create_regression, with which users can provide an R-style formula to automatically create the design matrix using the patsy Python package. Another method, specify_model, allows the specification of the desired parameters and dimensions to return. This method is used by create_inference to convert CmdStanPy output to ArviZ^58^ InferenceData objects.

There are two base classes included with BIRDMAn termed the TableModel and the SingleFeatureModel. The TableModel allows fitting an entire dataset at once, while the SingleFeatureModel allows for fitting individual features. The SingleFeatureModel is advantageous as it allows for highly parallelized workflows. Because there are often hundreds or thousands of features in a microbiome dataset, we note that using multiple CPUs to run many features at once is often more efficient than fitting the entire table. We provide a convenience class, ModelIterator, to iterate through the features in a given table. This class also allows for dividing the table into chunks. This allows users to customize the number of features to fit at once depending on their computational resources.

### Simulations

All simulations were performed through the fixed_param option in CmdStanPy. Ground-truth parameters were provided into a negative binomial generative model to simulate data from mean and dispersion parameters.

For the data-driven simulation, we randomly drew values for batch offset, batch dispersion, and base dispersion parameters. These parameters were fed into a model with ***β*_0_ = *N*(—8.1), *β*_1_ = *N*(2,1)**. Log sampling depth was simulated from a Poisson-Lognormal distribution with ***λ*** drawn from ***N***(5000, 0.2) We simulated 300 samples comprising 10 total batches with 10 total features.

For the variable sample size simulations, we simulated feature counts for 500 samples with ***β_0_ = 8, β_1_ = 3***, and 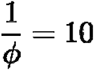. Log sequencing depths were simulated using a Poisson-Lognormal model with λ drawn from ***N***(50000,0.5) where depth varied.

To simulate variable rarefaction depth, we first drew ground truth intercept and beta values from ***N*(—8,1)** and ***N*(2,1)** respectively for 1000 features. These values were used to generate counts for 300 samples through the multinomial distribution. We used the multinomial distribution to enforce the same sampling depth for all samples, simulating rarefaction.

### Antibiotics case study

16S data was downloaded from Qiita study 494; we used 16S OTUs picked against the GreenGenes_13.8^59^ reference database at 97% sequence similarity. OTU picking was performed with SortMeRNA^60^ with Qiita default parameter values. Features present in fewer than 10 samples were filtered. We also removed samples with a total sequencing depth less than 1000.

To account for the longitudinal nature of this design, we used backwards difference encoding such that each time point was compared to the one immediately before it. We implemented the subject identifiers as a random effect with both random intercepts and random slopes. The posterior draws were centered around the mean. Ranking of OTUs by differentials for log-ratio feature selection was done using the posterior means.

We performed t-tests comparing the log-ratios between groups of samples at different timepoints. The alternative hypothesis was chosen such that samples from the later time point would have higher log-ratios than those from the initial timepoint due to the anticipated effect of Cp on microbial populations.

We then implemented multinomial logistic regression, random forest classification, and repeated k-fold cross-validation through scikit-learn for our machine learning approach. Because DESeq2 supports contrasts natively, we computed the same contrasts as BIRDMAn for parity. With ALDEx2 and ANCOM-BC, we computed the differentials associated with each timepoint using preCp as reference. For the random forest classifier, we used the CLR-transformed feature table (with a pseudocount of 1) entries as the predictors. All models were also provided one-hot-encoded vectors for subject identifiers. Performance was measured using balanced accuracy. For multinomial logistic regression we used the lbfgs solver with 1000 max iterations. For the random forest classifier we used a set random seed and 100 estimators. We used repeated stratified k-fold cross validation with 5 splits and 10 repeats and a random seed. All other parameters not mentioned were set to the scikit-learn defaults.

Posterior draws for timepoint-contrast differentials were analyzed with (1) FirstCp-associated features with preCp-associated features as reference and (2) SecondCp-associated features with Interim-associated features as reference. In this way, the posterior distribution reflects how each Cp course affects the selected bacterial features over time.

For determining the Gram status of each OTU, we used the BugBase^61^ web interface. We took the set intersection of Gram positive and Gram negative features with the features associated with both FirstCp and SecondCp to determine the Gram status breakdown of both numerator and denominator features.

### TCGA case study

The bacterial TCGA tables were obtained from those processed in Narunsky-Haziza et al.^62^ and Poore et al.^4^ All TCGA sequence data were accessed via the Cancer Genomics Cloud^63^ (CGC) as sponsored by SevenBridges (https://cgc.sbgenomics.com) after obtaining data access from the TCGA Data Access Committee through dbGaP (https://dbgap.ncbi.nlm.nih.gov/aa/wga.cgi?page=login). On Qiita^64^, TCGA WGS host-depleted and quality-controlled fastq files were used to generate a metagenomic table by direct genome alignments based on Woltka v0.1.1^65^ against the RefSeq^66^ release 200 (built as of May 14, 2020). The resulting tables can be found on Qiita under study ID 13722, of which we filtered to only analyze the bacteria and then were subsequently decontaminated through decontam^67^ (https://github.com/benjjneb/decontam) (version 1.14.0) following the protocol described in Poore et al.^4^

After initial table generation, we removed samples from data submitting centers with very few samples. We also filtered our data to only include samples from white, African-American, and Asian races. Additionally, we only included samples from patients who were alive at the time of sample procurement and retained only one sample per subject. To filter out lowly prevalent features, we removed features present in fewer than 50 total samples. To remove samples with low sequencing depth, we set a threshold of 500 reads. Finally, we included only cancer types with at least 20 instances in the dataset for statistical power.

We then built statistical models to model the differential associated with each cancer type. Because TCGA did not include “normal” samples from healthy individuals, we used breast cancer (BRCA) tumor samples as reference. Both race^68^ and gender were also included as covariates. Data submitting center was incorporated as a random effect (both random intercepts and random slopes).

Posterior means were computed for each feature’s association with each individual cancer type. For each cancer type, we ranked the differentials and used the top and bottom 200 features associated with that cancer type to compute log-ratios per sample. These log-ratios were used as predictor variables in our machine learning models.

Because not every cancer type was represented in each center, we performed multi-class classification within centers. For each center, we fit a model to predict cancer type from our log-ratios. This procedure was performed with 5 repeats of stratified 2-fold cross-validation. We repeated this machine learning process for cancer type differentials from DESeq2, ALDEx2, and ANCOM-BC. For comparison, we fit a random forest classifier on the CLR-transformed feature table to predict cancer type as well.

The leave-one-center-out models were fit using binomial logistic regression with balanced class weights. For each cancer type, we fit a model on all but one center and used that model to predict cancer type for the held-out center. We also used the same random forest classifier as previously described for comparison.

### Analysis & visualization software

Analysis of the results in this work were primarily performed through Python (v3.8.13). Pandas^69^ (v1.1.5) and NumPy^70^ (v1.22.3) were used for general data analysis. SciPy^71^ (v1.7.3) was used for computing statistical tests. For interfacing with multidimensional arrays we used xarray^72^ (v0.20.1) and ArviZ^58^ (0.12.1). Machine learning models were fit and cross-validated using scikit-learn^73^ (v1.0.2). Python figures were generated using seaborn^74^ (v0.11.2) and Matplotlib^75^ (v3.5.1) as well as Matplotlib-venn (v0.11.7). We used biom-format^57^ (2.1.12) and scikit-bio (v0.5.6) for statistical analysis of microbiome data structures.

R analysis was performed using the tidyverse^76^ packages dplyr (v1.0.9), stringr (v1.4.0), and ggplot2 (v3.3.6). Phylogenetic visualization was performed using treeio^77^ (v1.18.0) and ggtree^78^ (v3.2.0). BIOM tables were read using the biomformat R package (v1.22.0).

## Code and data availability

All data used were downloaded from publicly available Qiita studies. The scripts and Stan models used to analyze these data as well as Jupyter notebooks for the visualizations are available at https://github.com/knightlab-analyses/birdman-analyses-final. The BIRDMAn software package is available at https://github.com/biocore/BIRDMAn and the documentation is available at https://birdman.readthedocs.io/. All analyses in this work were performed using BIRDMAn v0.1.0.

## Acknowledgments

We thank the members of the Knight Lab and Morton Lab for feedback and bug reporting for the BIRDMAn software. We thank the developers of ArviZ and CmdStanPy for responding to and addressing issues on GitHub as well as the users on the Stan forums for answering our questions.

This work was supported in part by NIH U19AG063744, NIH 1DP1AT010885, and NIH U24CA248454. J.T.M. was funded by the intramural research program of the Eunice Kennedy Shriver National Institute of Child Health and Human Development (NICHD).

## Author information

G.R., J.T.M., and R.K. conceived the idea for the study. G.R. & J.T.M. developed the BIRDMAn software package. G.R., J.T.M., C.G., G.D.S-P., & C.M. contributed to the case study and simulation analysis. C.A., J.T.M., C.M., & R.K. helped to define the scope of the analyses. G.R. & Y.C. contributed to the documentation for BIRDMAn. M.E., Y.C., D.H., & C.M. gave critical feedback on the usage and documentation of the software. All authors helped write and review the manuscript.

## Supplementary Figures

**Supplementary Fig 1:**
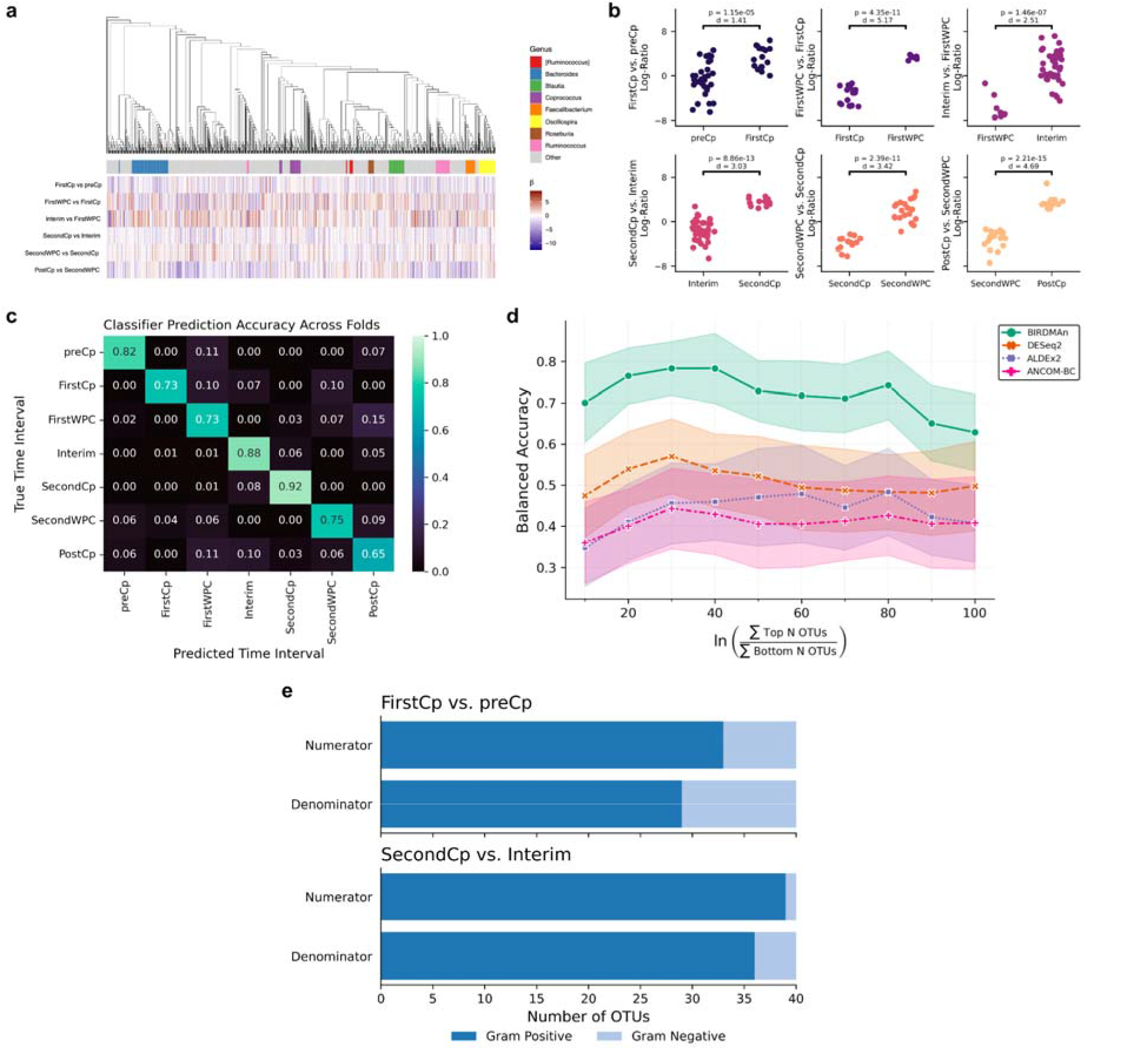
(a) Phylogenetic tree of all OTUs with a heatmap of posterior means for each time-interval contrast. OTUs assigned to one of the top 8 most abundant genera are annotated through the colored strip. (b) When BIRDMAn is used to account for per-subject variation, log-ratio comparisons of the top 40 OTUs vs. bottom OTUs are associated with the difference between each time point and the next one. For each of these contrasts, the log-ratios of the samples between the two time intervals were compared using a one-sided t-test. Plots are annotated with p-values. Different taxa contribute to the log ratios for each contrast. (c) Overall performance of BIRDMAn classifier on predicting the antibiotics time interval using the log-ratios. The classifier prediction accuracies shown are aggregated across folds and repeats from repeated k-fold cross-validation. (d) Accuracy of the multinomial classifier by number of OTUs used in log-ratio calculations. Points represent mean accuracy across cross-validation iterations and shaded areas represent ±1 standard deviation. (e) Distribution of Gram positive and Gram negative OTUs associated with FirstCp and SecondCp log-ratios.

**Supplementary Fig 2:**
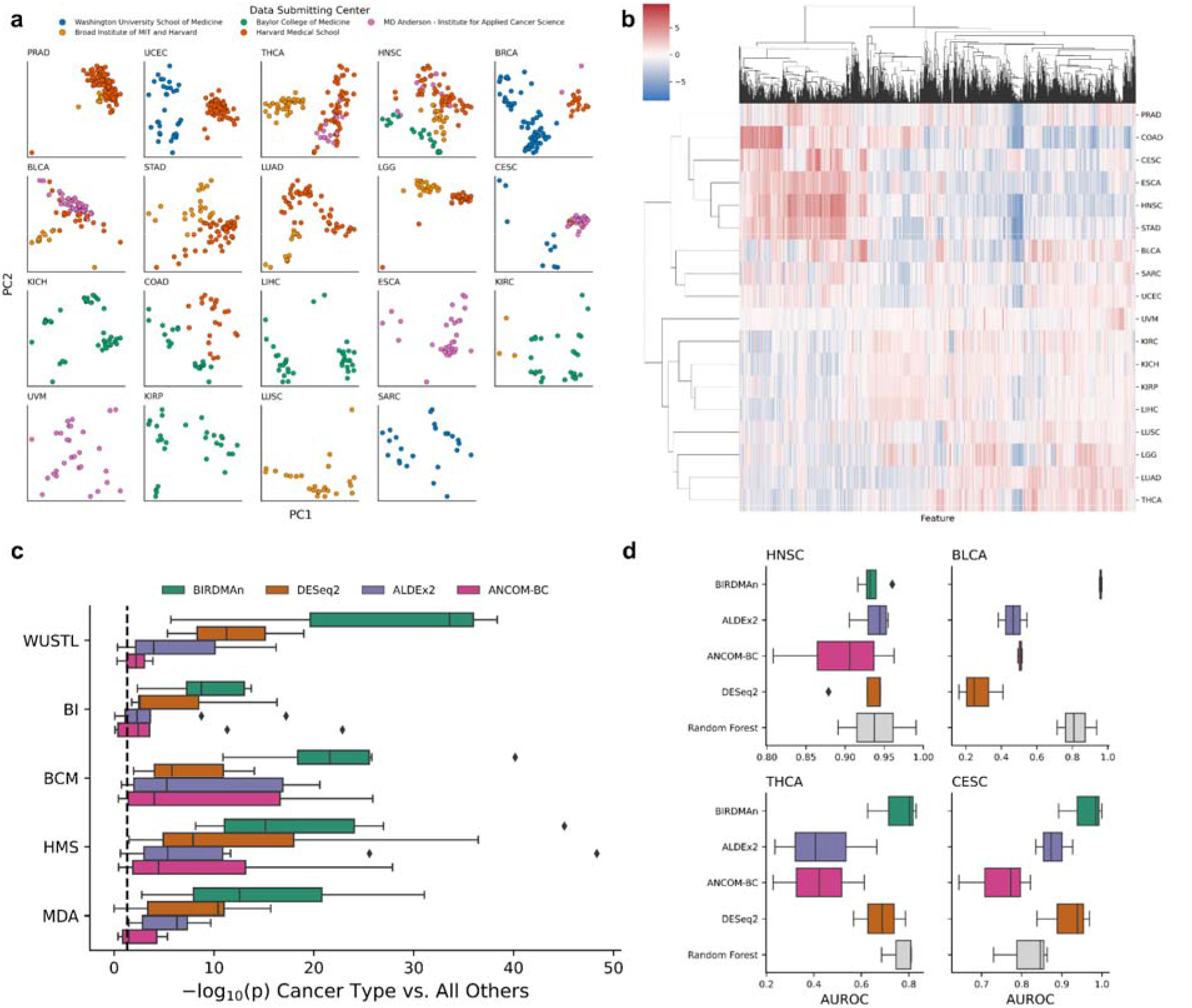
(a) RPCA projection of the original feature table subset to each individual cancer type. Points are colored by data submitting centers, showing that many cancer types exhibit strong separation by batch. (b) Posterior means (CLR) of feature differentials clustered by cancer type. (c) Log-ratios identified by BIRDMAn separate each tumor type from all others when stratified by center. Dashed line represents a t-test p-value at p = 0.05. (d) Performance of leave-one-center-out cross-validation logistic regression classifier AUROC of all methods.

